# Urinary Exosomal miRNA Profiling Reveals Sensitive Non-Invasive Diagnosis of Bladder Cancer

**DOI:** 10.1101/2025.10.09.681526

**Authors:** Garima Singh, Anil Kumar, Lalit Kumar, Neelu Mishra, Shristy Bhattacharjee, Kunal Yadav, Yashasvi Singh, Ujwal Kumar, Sameer Trivedi, Samarendra K. Singh

## Abstract

**Unstructured:** miRNAs represent a transformative advancement in both research and clinical management of urinary bladder cancer (UBC), emerging as clinically significant molecular targets with the potential to revolutionize existing diagnostic standards. This study exploited the robust stability of miRNAs and profiled urinary miRNAs in UBC patients and controls through miRNA sequencing, revealing an expanded and altered miRNA repertoire in cancer samples. Validation in an independent cohort revealed fold changes for miR-6724-5p, miR-1273h-5p, miR-7704, miR-200-5p, and miR-10400-5p ranged from ∼46 to ∼2,777 (p < 0.01). Stage-specific analyses highlighted dynamic miRNA expression linked to tumor progression. Diagnostic evaluation identified an optimal three-miRNA panel comprising miR-6724-5p, miR-10400-5p, and miR-7704, achieving diagnostic accuracies (AUC > 70%) and high sensitivity (>90%). These data demonstrate the potential of urinary mature miRNAs as robust biomarkers for non-invasive detection and monitoring of UBC in early stages, supporting their translation into clinical diagnostic assays pending larger prospective studies.

**Structured:** *Background:* Urinary microRNAs (miRNAs) are promising candidates due to their molecular stability and disease-specific expression. miRNAs represent a transformative advancement in both research and clinical management of urinary bladder cancer (UBC), emerging as actionable molecular targets with the potential to revolutionize existing diagnostic standards.

*Methods:* Small RNA sequencing was performed on urine samples from UBC patients and healthy controls to profile miRNA expression. Mature miRNAs were prioritized by exclusion of precursor forms using computational filtering. Validation by quantitative RT-PCR was conducted on an independent, age-matched cohort (n = 45). Diagnostic potential of candidate miRNAs was assessed by receiver operating characteristic (ROC) curves and area under the curve (AUC) analyses.

*Results:* Sequencing identified 865 known and 11 novel miRNAs, with UBC samples showing greater miRNA diversity (708 known miRNAs) compared to controls (540). Ten miRNAs were significantly dysregulated in UBC (p < 0.05). Validation revealed fold changes for miR-6724-5p, miR-1273h-5p, miR-7704, miR-200-5p, and miR-10400-5p ranged from ∼46 to ∼2,777 (p < 0.01). Stage-specific analyses highlighted dynamic miRNA expression linked to tumor progression. A three-miRNA panel (miR-6724-5p, miR-10400-5p, miR-7704) demonstrated superior diagnostic performance (AUC > 70%, sensitivity > 90%).

*Conclusion:* Mature urinary miRNAs exhibit reproducible, stage-specific dysregulation in UBC and a combinatorial panel offers sensitive, specific non-invasive detection. These findings warrant further multicenter validation for clinical application.

## Introduction

Urinary bladder cancer (UBC), one of the most widespread malignancies, stands as the 9^th^ most commonly diagnosed cancer worldwide, with rising mortality rates in recent years. In 2022 over 614,298 new bladder cancer cases were reported globally, representing a 7.1% increase compared to 2020 and ranking it 13^th^ in terms of mortality(Bray et al., 2024). The burden of bladder cancer is approximately three to four times higher in men than in women, with rates rising steeply among older adults, with risk factors including smoking, occupational chemical exposure, and chronic bladder inflammation and urinary tract infections (UTIs), showcasing symptoms such as pelvic pain, haematuria, and frequent urination (Chesnut et al., 2021). Of all cases of urinary bladder cancer reported, urothelial carcinoma constitutes over 90% of the cases, accounting for the most frequent cases (Heath & Rosenberg, 2021). In regions such as India, where industrialization, environmental pollution, and tobacco usage are on the rise, the incidence of urinary bladder cancer has escalated alarmingly, contributing to ∼3.9% of all cancer diagnoses nationwide, with an age-adjusted incidence rate of 3.7 to 4.0 per 100,000 among males and 0.8 to 1.3 per 100,000 among females showing wide variation between regions (Kumar et al., 2022; Mishra & Balasubramaniam, 2020; Zhang et al., 2022). Despite advances in therapeutic strategies, UBC outcomes remain suboptimal, primarily due to delayed detection and high recurrence rates (50-90% of the total cases)(Kamat et al., 2017). The current gold standards for diagnosis and surveillance are cystoscopy and urine cytology. However, cystoscopy is limited by invasiveness and patient discomfort whereas urine cytology is limited by poor sensitivity for low-grade tumors (Oeyen et al., 2019; Zhu et al., 2019). These constraints highlight an urgent need for robust, non-invasive and sensitive biomarkers to facilitate early detection and improve clinical management.

MicroRNAs (miRNAs) have emerged as promising biomarkers in this context due to their stability in urine and their ability to reflect disease-specific molecular changes in bladder cancer (Chattopadhaya et al., 2025; Miah et al., 2012; Urquidi et al., 2016). miRNAs are a class of small, noncoding RNAs that post-transcriptionally regulate gene expression and play pivotal roles in cellular functions. Aberrant expression of miRNAs has been implicated in the initiation and progression of various malignancies, including UBC(Arenas et al., 2022; Bañuelos-Villegas et al., 2021; Feng et al., 2015; Miah et al., 2012; Tripathy et al., 2025). Distinct urinary miRNA signatures can differentiate cancerous from non-cancerous states and also discern disease stages with diagnostic accuracy comparable or superior to traditional protein markers (Sapre et al., 2016). Thus, exploring the dysregulated miRNAs in Urinary bladder cancer patients could unfold new and amended diagnostic approaches, significantly differing from previously existing, expensive, and highly invasive procedures.

Importantly, miRNAs are released into circulation and other body fluids, including urine, highlighting their impact on survival and tumor communication. Their remarkable stability against enzymatic degradation enables consistent and reliable detection (Guhaniyogi & Brewer, 2001; Valadi et al., 2007). Coupled with their cancer-specific expression patterns, miRNAs potentiate as robust non-invasive biomarkers for diagnosis, prognosis, and therapeutic monitoring in urinary bladder cancer. Exploring miRNA signatures unique to bladder cancer may therefore offers critical insights into tumor biology while paving the way for the development of more sensitive, easily accessible, and cost-effective miRNA-based diagnostic panels or algorithms for improved detection and monitoring of bladder cancer.

In this study, we conducted a comprehensive and systematic screening of urinary miRNA profile using high-throughput sequencing, followed by validation of differentially expressed miRNAs through RT-qPCR assays. Seven mature miRNAs exhibiting significant differential expression were systematically characterized in relation to UBC. Also, the study correlates the clinical parameters and associates the particular miRNA with the different stages of urinary bladder cancer. Moreover, our findings lay a foundation of urine miRNAs as potential biomarkers for early-stage diagnosis and provide a step towards effective and precise surveillance of UBC. By integrating molecular data with clinical outcomes, we aim to define robust, non-invasive biomarkers that enable early detection, inform treatment decisions, and improve prognostic accuracy, ultimately improving survival and quality of life for urinary bladder cancer patients.

## Methodology

### 1. Study Design

This was a single-centre, observational study conducted at the, School of Biotechnology, Institute of Science, in collaboration with the Department of Urology, Institute of Medical Sciences (IMS) Banaras Hindu University (BHU), Varanasi, Uttar Pradesh. A total of 48 participants (32 patients, 16 controls) were selected during the study from March 2023 to April 2025. The study protocol and the patient consent procedures were approved by the Ethical Committee, IMS-BHU (Ref. No. 2024/EC/7064). Written informed consent was obtained from all participants (in both Hindi/ English) after a detailed explanation of the study protocol.

### 2. Study Population and Sampling

#### 2.1 Inclusion and exclusion Criteria

Participants’ data were collected from the Outpatient Department (OPD) of the Department of Urology at Sir Sunderlal Hospital (SSH), IMS-BHU.

**Inclusion**: Individuals had a histopathologically confirmed, newly diagnosed case of UBC with no prior treatment; were aged between 18 and 80 years; had an Eastern Cooperative Oncology Group (ECOG) performance status score ranging from 0 to 2; and demonstrated a willingness to participate in the study, regardless of gender.

**Exclusion:** Individuals were excluded from the study if they had a prior history of bladder malignancy, a concurrent diagnosis of other systemic cancers, or had previously received chemotherapy or radiotherapy. Additionally, individuals with an active UTI, as well as those who declined or were unwilling to provide informed consent, were not considered eligible for participation.

#### 2.2. Sampling

Based on the historical case inflow, an estimated incidence of 3 cases/month was anticipated, yielding approximately 32 patients over the study period. For each case, an age-matched healthy control was recruited in a 2:1 ratio, resulting in a final cohort of 32 cases and 16 matched controls. (Table 1).

**Table 1:** Summary of participants studied in the report.

| Gender | Controls |  | UBC Cases |  |
| --- | --- | --- | --- | --- |
|  | n | % | n | % |
| Male | 13 | 81.3 | 27 | 84.4 |
| Female | 3 | 18.7 | 5 | 15.6 |
| Age |  |  |  |  |
| Median | 69 |  | 68 |  |
| Average | 66 |  | 65 |  |
| Range | 44-78 |  | 44-78 |  |
| EAU Risk Classification |  |  |  |  |
| Low |  |  | 5 |  |
| Intermediate |  |  | 14 |  |
| High |  |  | 5 |  |
| MIBC |  |  | 5 |  |
| Metastatic |  |  | 3 |  |
| HPE Stage |  |  |  |  |
| Ta |  |  | 10 |  |
| T1 |  |  | 16 |  |
| T2 |  |  | 6 |  |
| Total | 16 |  | 32 |  |

Abbreviations: UBC = urinary bladder cancer. n = total number. % = percentage. EAU= European Association of Urology. HPE= histopathological examination. MIBC = muscle invasive bladder cancer. Ta, T1, T2 = different stages of tumor.

All patients underwent a comprehensive pre-operative evaluation that encompassed clinical, laboratory, and radiological assessments. A detailed clinical history and physical examination were conducted to assess the general health status and identify any signs indicative of disease progression. Laboratory investigations included complete blood count (CBC), kidney function tests (KFT), liver function tests (LFT), serum calcium levels, random blood sugar (RBS) levels, and urinalysis comprising routine examination, culture and sensitivity testing, and cytological analysis. Radiological evaluation involved multiparametric magnetic resonance imaging (mpMRI) of the pelvis following the VI-RADS protocol to determine local tumour staging, along with either non-contrast computed tomography (NCCT) or high-resolution computed tomography (HRCT) of the thorax to assess for metastatic spread. Following treatment, patients were monitored at 3-month and 6-month intervals to evaluate clinical progression and survival outcomes.

### 3. Sample Collection and RNA Extraction

Freshly voided 5 ml urine samples were collected from 48 participants prior to instrumentation/ bladder tumor removal. Urine was stored at 4 °C for 2-4 hours before freezing at -80 °C to prevent degradation. Samples were subjected to thawing followed by RNA isolation using the mirVana kit (Ambion, Austin,TX, USA) according to the manufacturer’s protocol and quantified using Qubit RNA BR Assay.

### 4. Library Preparation Protocol

100 ng of RNA was used for library preparation following the QIAseq miRNA Library Kit protocol, which involves ligating adapters sequentially to the 3’ and 5’ ends of miRNAs. The ligated miRNAs were then converted to cDNA by using a reverse transcription primer with Unique Molecular Identifier (UMI). The library was proceeded for cleanup and amplified using the following thermal conditions: Hold 95°C for 15 mins; 16 cycles of 95°C for 15sec, 60°C for 30sec, 72°C for 15sec; Final extension of 72°C for 2mins and hold 4 °C for hold. The amplified libraries were further cleaned up using a magnetic bead-based method and checked for fragment size distribution on TapeStation using High Sensitivity D1000 ScreenTape (5067-5584).

### 5. miRNA Sequencing and analysis

Cluster amplification of the library was then carried out on an Illumina flow cell, and sequencing was performed using Illumina NextSeq 550. Raw sequencing data obtained in FASTQ format were initially demultiplexed to separate the samples. An initial quality assessment and pre-processing of the sequencing reads were performed using FastQC (http://www.bioinformatics.babraham.ac.uk/projects/fastqc). Based on the results from the FastQC quality reports, low-quality portion of the reads was trimmed.

The raw sequencing data were further processed to trim low-quality bases from the 3’ ends of reads using the cutadapt tool. Reads were then length-filtered to select sequences between 17 and 30 nucleotides, which are appropriate for small RNA analysis. The collapsed reads were aligned to miRbase reference sequence using Bowtie software with default parameters. Alignment files were further processed for editing and indexing with Samtools, and Bedtools was employed to assess overlaps between aligned reads and the genome. Known miRNAs were profiled by mapping reads to whole miRNA sequences in the miRbase database. Further, the sub-sequences data were retrieved from FASTA files using the faidx tool. For the discovery and quantification of novel miRNAs, miRDeep2 was utilized. The expression values of known miRNA, determined using DESeq2, were used for differential expression analysis. The significant miRNA was selected based on p-value < 0.05. Heatmap and DE plot for significantly expressed miRNA were plotted using gplots, iDEP 2.1 and SR Plot.

### 6. Quantitative Real-Time PCR (qRT-PCR)

Total RNA extracted (500 ng to 1 µg) was used for first-strand complementary DNA (cDNA) synthesis employing the PrimeScript™ RT Reagent Kit (Takara Bio, Japan) according to the manufacturer’s protocol. Stem-loop primers specific to the target miRNAs (Table 2) were used during reverse transcription. The generated cDNA was subsequently used as a template for quantitative real-time PCR, performed with TB Green® Premix Ex Taq™ II (Takara Bio, Japan) using miRNA-specific primer sequences (Table 2) on an Applied Biosystems QuantStudio 5 system.

**Table 2:**
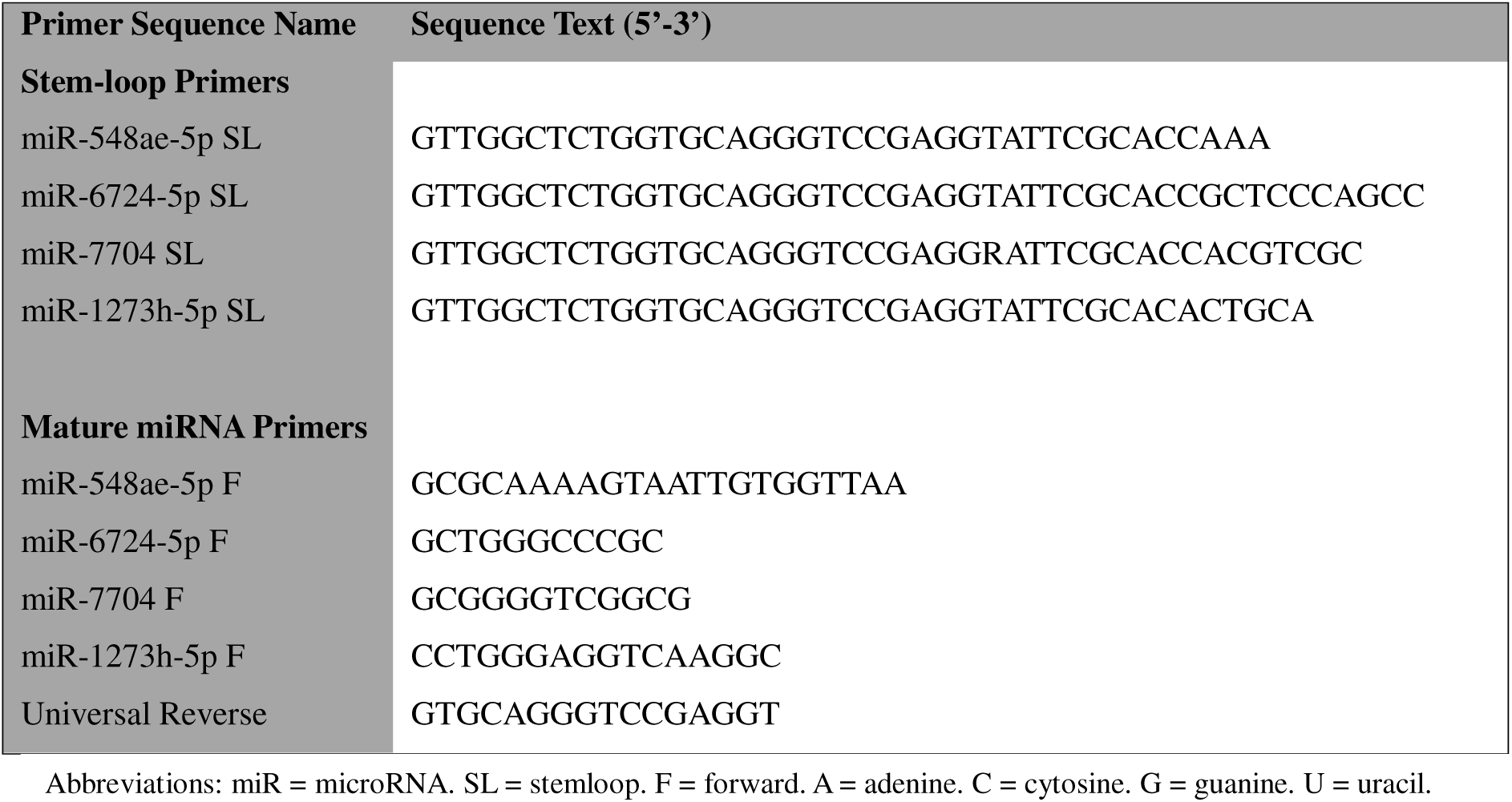
List of Primer Sequences.

### 7. Statistical Analysis

All the continuous data were expressed as mean ± standard deviation. Relative miRNA concentration was calculated with respect to reference U6 (ΔCt=Ct miR-Ct U6) and the fold change was calculated using the 2^-ΔΔCt^ function (Schmittgen & Livak, 2008). Comparison was performed using T-test with unequal variance. Descriptive statistics, including mean, standard deviation, and percentages, were calculated to summarize demographic and clinical characteristics of the study population. For variables that did not follow a normal distribution, non-parametric tests were employed: the Mann–Whitney U-test was used for comparisons between two independent groups. A p-value of ≤ 0.05 was considered statistically significant throughout the analysis.

## Results

### miRNA profiling reveals an altered urinary miRNA landscape in bladder cancer

High-throughput miRNA sequencing of urine samples from a control and UBC patient yielded over 18 million and 54 million reads, respectively, with high base quality and a predominant read length around 22 nucleotides consistent with mature miRNAs (Supplementary Figure 1A and 1B). Mapping to miRBase detected 865 known miRNAs along with 11 novel candidates across both samples. Compared to control urine, the UBC sample exhibited a more complex profile with 708 known miRNAs versus 540 in the control, suggesting cancer-associated miRNA dysregulation (Figure 1A and 1B). Further analysis for differential expression identified 10 significantly dysregulated known miRNAs (p < 0.05; defined by thresholds of log2Foldchange ≥ 1.0 for upregulation and ≤ –1.0 for downregulation), comprising one upregulated and nine downregulated miRNAs in the cancer sample (Figure 1C). Since precursor RNA are not biologically active and may not accurately reflect disease-specific changes in urine therefore, stringent filtering was performed to exclude precursor sequences, prioritized mature miRNAs for further validation and functional interpretation (Figure 1D and Supplementary Table 1).

**Figure 1:**
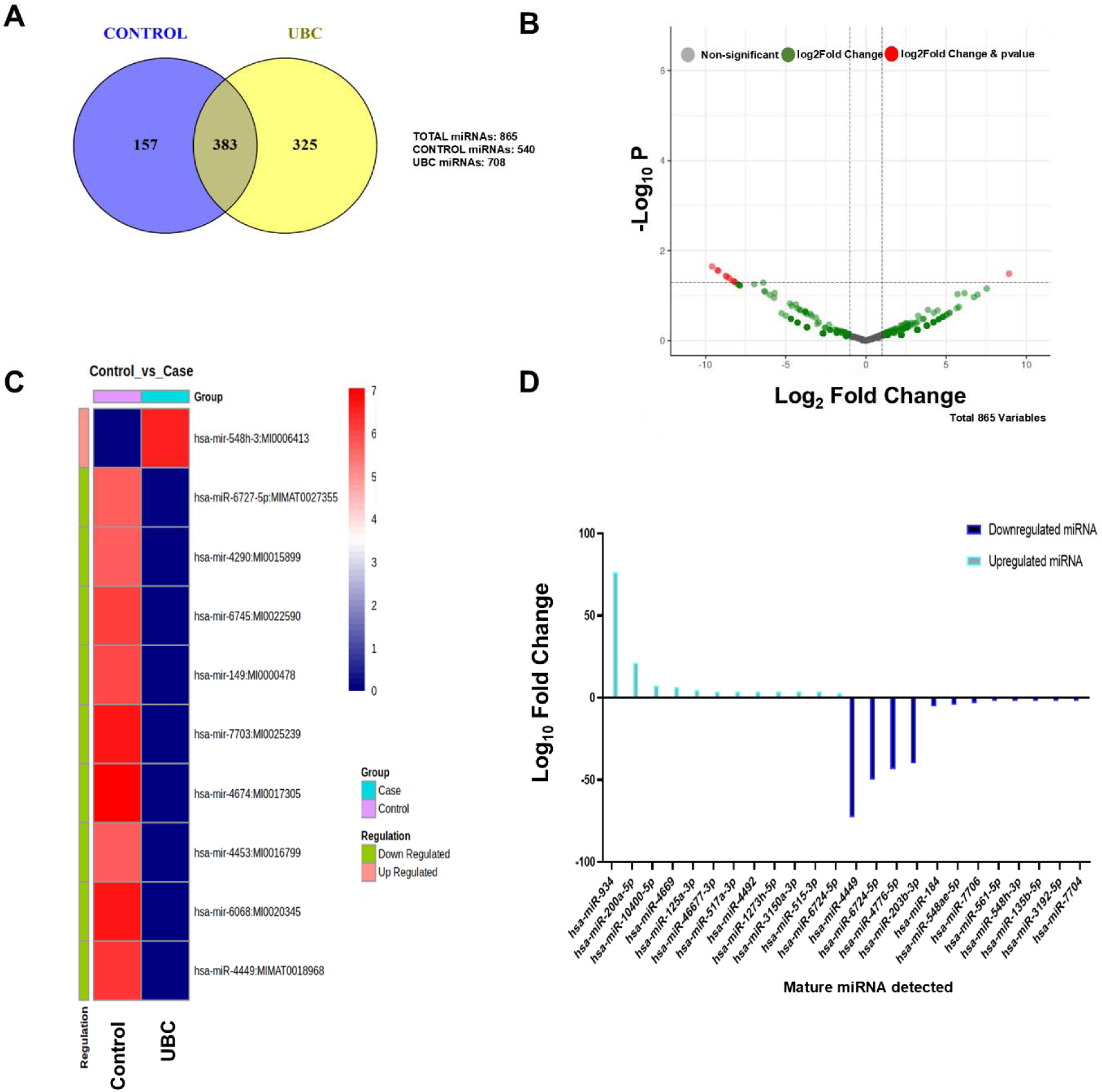
Analysis of differentially expressed microRNAs. **A.** Venn diagram of total 865 miRNAs identified in Urinary Bladder Cancer Patient and Control samples, among which 325 miRNAs are present exclusively in UBC patient samples, 157 miRNAs in control samples, and 383 miRNAs in both the samples. **B.** Volcano plot showing the differentially expressed miRNAs. x-axis represents log2 fold change, y-axis represents statistically significant values (padj= adjusted p-value, threshold < 0.05). **C.** Heat map showing the varying expression of 10 miRNAs with most significant differential expression in control and UBC patient group, where, deep-red and deep-blue indicates the most significant upregulation and downregulation respectively. **D.** Bar plot showing the relative expression levels of the significantly expressed mature miRNAs, where dark blue and light blue indicate the downregulated and upregulated miRNAs, respectively.

### Validation in expanded cohort highlights heterogeneous miRNA expression and robust miRNA biomarker candidates

To identify miRNAs with potential as biomarkers, and prepare an unbiased biomarker panel for UBC detection, the sequencing data was validated on an expanded cohort of age-matched 45 participants for seven miRNAs. Among which four of them were found to be exclusively expressed in UBC cases (miR-200a-5p, miR-10400-5p, miR-1273h-5p and miR-6724-5p), two of them were exclusively associated with control case (miR-548ae-5p and miR-7704) and one miRNA was associated both with case and control but was found to be upregulated in UBC (miR-934) (Supplementary Table 1). The expression of miRNAs was varied from ∼2 fold to 3000 fold from lowest (miR-934) to highest (miR-200-5p) (Table 3). We quantified the expression of miRNAs across patient samples and controls and observed a significant upregulation of miR-6724-5p with an average fold change of 1,494.09 in UBC samples compared to controls (*p* = 0.0008) (Figure 2B). Similarly, miR-1273h-5p and miR-200-5p exhibited robust overexpression, showing average fold changes of 1,085.86 (*p* = 0.0069) and 2,776.52 (*p* = 0.0011), respectively (Figure 2C and 2F). MiR-10400-5p displayed significant upregulation with a fold change of 45.88 (*p* = 0.0011) (Figure 2G).

**Figure 2:**
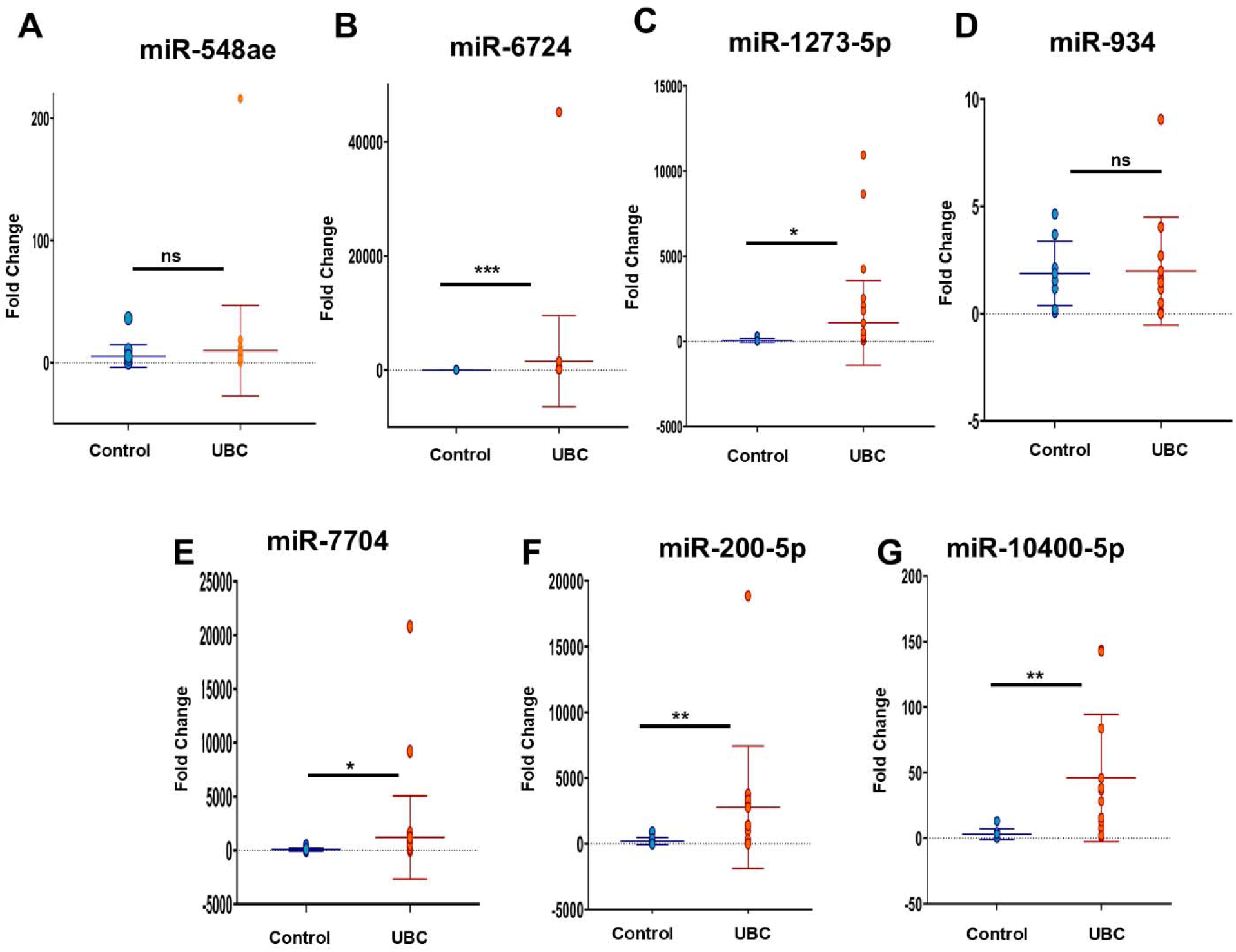
Validation of differentially expressed microRNAs across control and UBC patient samples. Aligned dot plots (lines at Mean ± SD) show the expression profiles of each microRNA according to the presence of bladder cancer. **A.** miR-548ae. **B.** miR-6724-5p. **C.** miR-1273-5p. **D.** miR-934. **E.** miR-7704. **F.** miR-200-5p. **G.** miR-10400-5p. (Abbreviations: * = p – value < 0.05. ** = p – value < 0.01. *** = p – value < 0.001. ns = not significant).

**Table 3:**
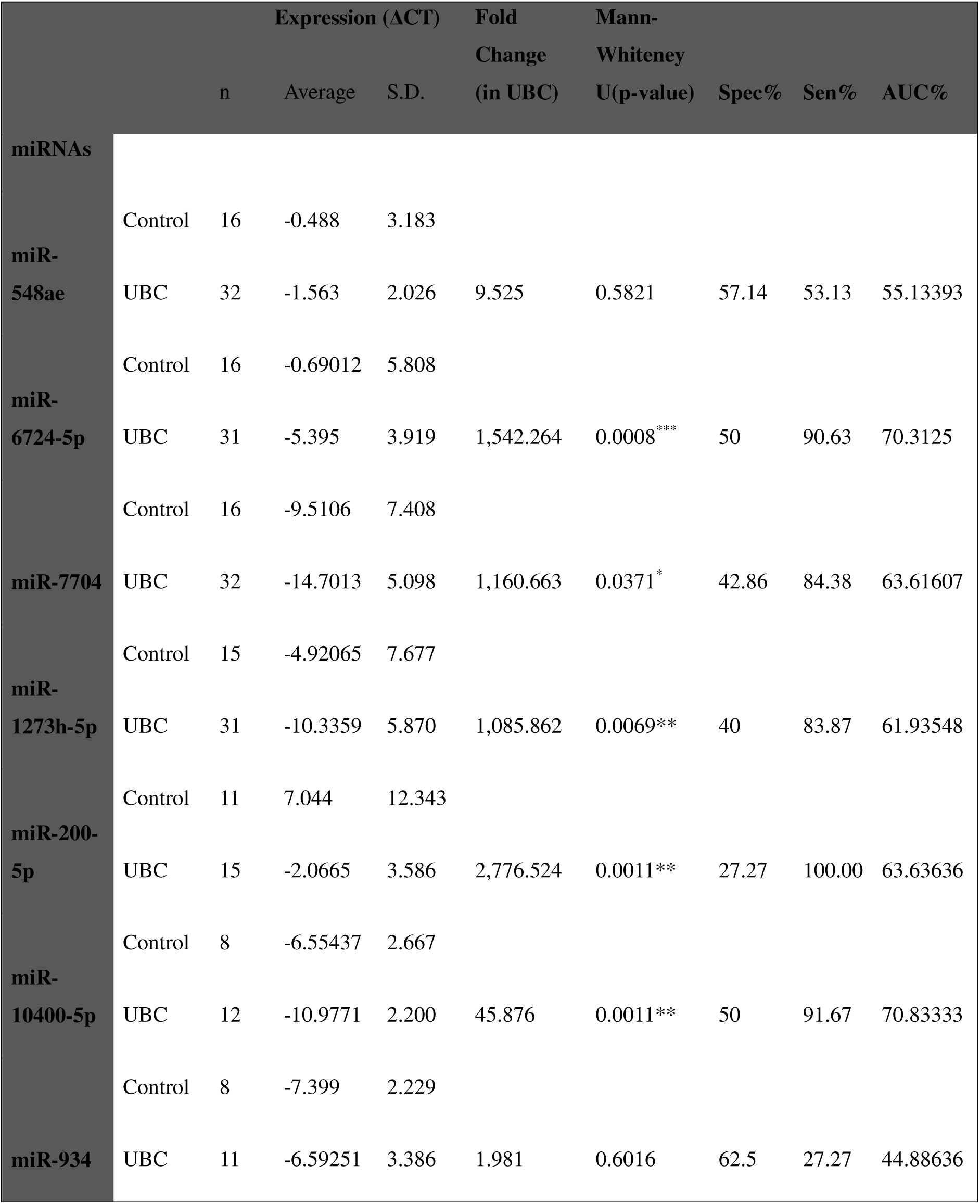
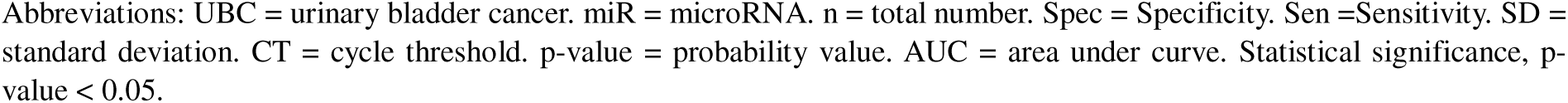
MicroRNA expression in urinary samples from patients with bladder cancer and controls.

|  |  | n | Expression (ΔCT) |  | Fold Change<br>(in UBC) | Mann-Whitney<br>U(p-value) | Spec% | Sen% | AUC% |
| --- | --- | --- | --- | --- | --- | --- | --- | --- | --- |
|  |  |  | Average | S.D. |  |  |  |  |  |
| miRNAs |  |  |  |  |  |  |  |  |  |
| miR-548ae | Control | 16 | -0.488 | 3.183 |  |  |  |  |  |
|  | UBC | 32 | -1.563 | 2.026 | 9.525 | 0.5821 | 57.14 | 53.13 | 55.13393 |
| miR-6724-5p | Control | 16 | -0.69012 | 5.808 |  |  |  |  |  |
|  | UBC | 31 | -5.395 | 3.919 | 1,542.264 | 0.0008*** | 50 | 90.63 | 70.3125 |
| miR-7704 | Control | 16 | -9.5106 | 7.408 |  |  |  |  |  |
|  | UBC | 32 | -14.7013 | 5.098 | 1,160.663 | 0.0371* | 42.86 | 84.38 | 63.61607 |
| miR-1273h-5p | Control | 15 | -4.92065 | 7.677 |  |  |  |  |  |
|  | UBC | 31 | -10.3359 | 5.870 | 1,085.862 | 0.0069** | 40 | 83.87 | 61.93548 |
| miR-200-5p | Control | 11 | 7.044 | 12.343 |  |  |  |  |  |
|  | UBC | 15 | -2.0665 | 3.586 | 2,776.524 | 0.0011** | 27.27 | 100.00 | 63.63636 |
| miR-10400-5p | Control | 8 | -6.55437 | 2.667 |  |  |  |  |  |
|  | UBC | 12 | -10.9771 | 2.200 | 45.876 | 0.0011** | 50 | 91.67 | 70.83333 |
| miR-934 | Control | 8 | -7.399 | 2.229 |  |  |  |  |  |
|  | UBC | 11 | -6.59251 | 3.386 | 1.981 | 0.6016 | 62.5 | 27.27 | 44.88636 |

Abbreviations: UBC = urinary bladder cancer. miR = microRNA. n = total number. Spec = Specificity. Sen =Sensitivity. SD = standard deviation. CT = cycle threshold. p-value = probability value. AUC = area under curve. Statistical significance, p-value < 0.05.

Interestingly, miR-7704 demonstrated a contrasting expression pattern upon validation. Whereas sequencing data indicated it as downregulated in UBC, qRT-PCR consistently showed its significant upregulation with an average fold change of 1,160.66 (*p* = 0.0371). This discrepancy underscores the importance of independent validation and suggests possible technical or biological factors influencing miR-7704 detection between platforms (Figure 2E).

MiR-548a-5p and miR-934 exhibited modest fold changes and did not achieve statistical significance (*p* > 0.5), indicating variable expression or a requirement for larger cohorts to clarify their roles. Collectively, these findings validate the majority of sequencing discoveries and reinforce the potential of these miRNAs as urinary biomarkers for UBC (Figure 2A and 2D).

### Stage-specific association of dysregulated key miRNAs in UBC revealed miR-200-5p maintained a high level from low to high risk cancer

To better understand the biomarker potential of the miRNAs across different stages of bladder cancer progression (including low-risk, intermediate, high-risk, muscle-invasive bladder cancer (MIBC), and metastatic stages) and clinical prognosis, we assessed the association between the miRNA expression patterns and clinical status in UBC patients. Several miRNAs exhibited significant upregulation during cancer progression, suggesting their potential stage-specific roles in tumor biology. miR-548ae showed moderate fold increases across all stages, peaking at intermediate and high-risk conditions (∼2.7–2.9 fold) but remained close to baseline in metastatic samples (∼1-fold), suggesting early involvement in tumor growth rather than metastasis (Figure 3A). In contrast, miR-6724-5p showed substantial upregulation, particularly in metastatic cases (225-fold), indicating its possible role in advanced cancer stage or aggressiveness (Figure 3B). miR-7704 displayed significantly elevated levels at the intermediate risk condition with >1000-fold increase, reflecting dynamic regulation during tumor development and potential as a progression marker (Figure 3E). Notably, miR-1273h-5p expression surged dramatically in metastatic samples (2740-fold), implying involvement in metastatic processes (Figure 3C). Among all the miRs, miR-200-5p maintained elevated levels in early to high-risk cases (2000–5600 fold) but was reduced in the metastatic stage (∼770 fold), pointing towards a critical role in tumorigenesis and possible loss of function during metastasis (Figure 3F). miR-10400-5p and miR-934 showed variable but lower expression changes, with miR-934 demonstrating reduced expression in high-risk and MIBC stages (Figure 3D and 3G).

**Figure 3:**
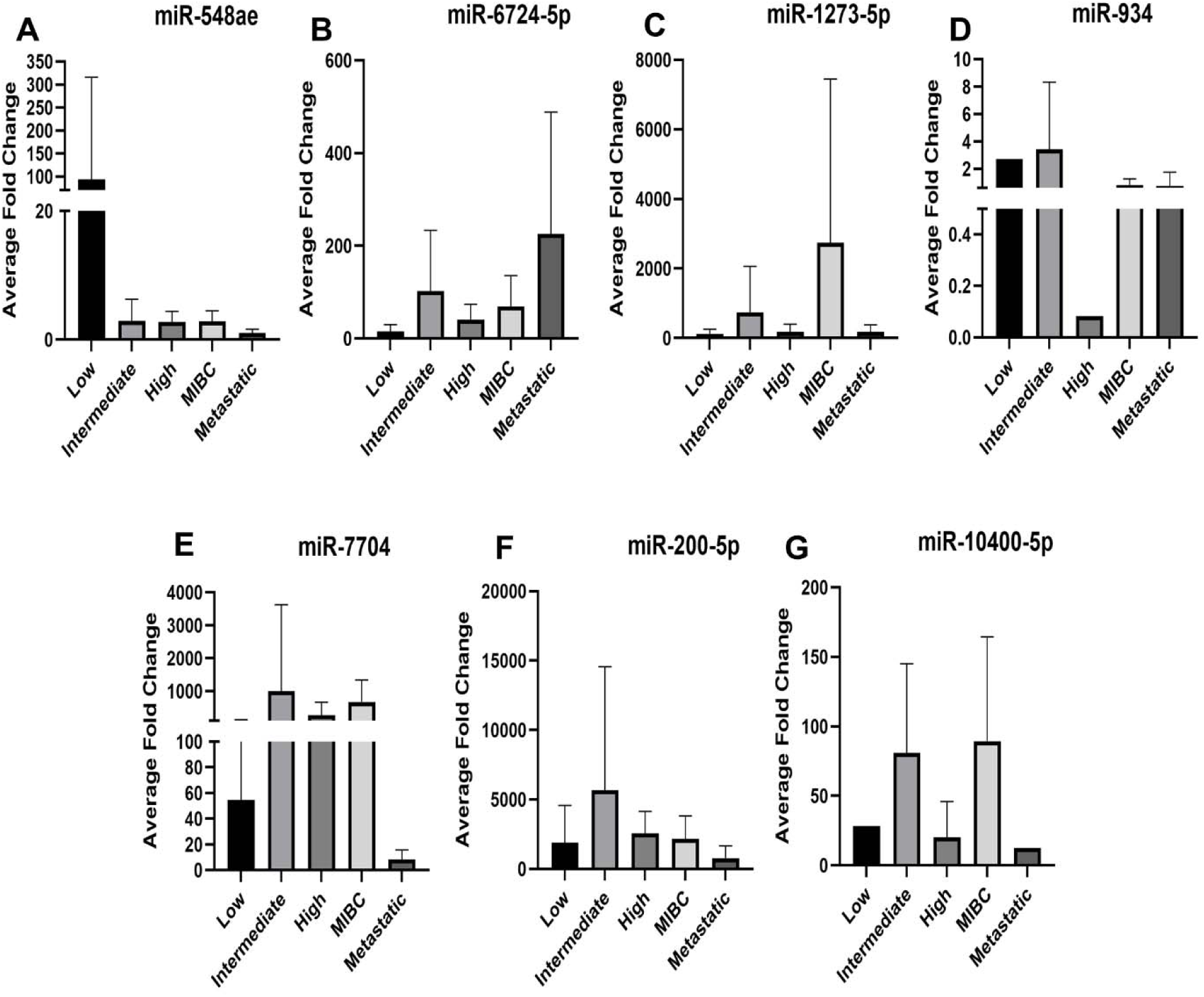
UBC risk/stage specific expression levels of the microRNAs. Box plots (Mean ± SD) showing differential expression of miRNAs in different stages of UBC; x-axis demonstrating the cancer progression risk/stage, y-axis showing the average foldchange (average 2^-ΔΔct^) for the particular miRNA. **A.** miR-548ae. **B.** miR6724-5p. **C.** miR-1273-5p. **D.** miR-934. **E.** miR-7704. **F.** miR-200-5p. **G.** miR-10400-5p.

### miR-6724/7704/10400 panel serves as a non-invasive diagnosis of bladder cancer

We further calculated the diagnostic performance of differentially expressed miRNAs for UBC, using the ΔCt value (Table 3). Quantitative evaluation of seven candidate miRNAs for urinary bladder cancer (UBC) diagnosis revealed variable diagnostic performances as reflected by specificity, sensitivity, and AUC values (see Table). The sensitivity of individual miRNAs varied from 27.27% to 100 % and the specificity varied from 27.27% to 62.50% in UBC. miR-6724-5p and miR-10400-5p yielded the highest AUC scores (70.31% and 70.83%, respectively) and exhibited robust sensitivity (>90%). miR-7704 and miR-200-5p displayed high sensitivities (84.38% and 100%, respectively) but showed moderate or lower specificity. Conversely, miR-934 demonstrated the highest specificity (62.5%) but limited sensitivity (27.27%), resulting in the lowest AUC (44.89%).

**Figure 4:**
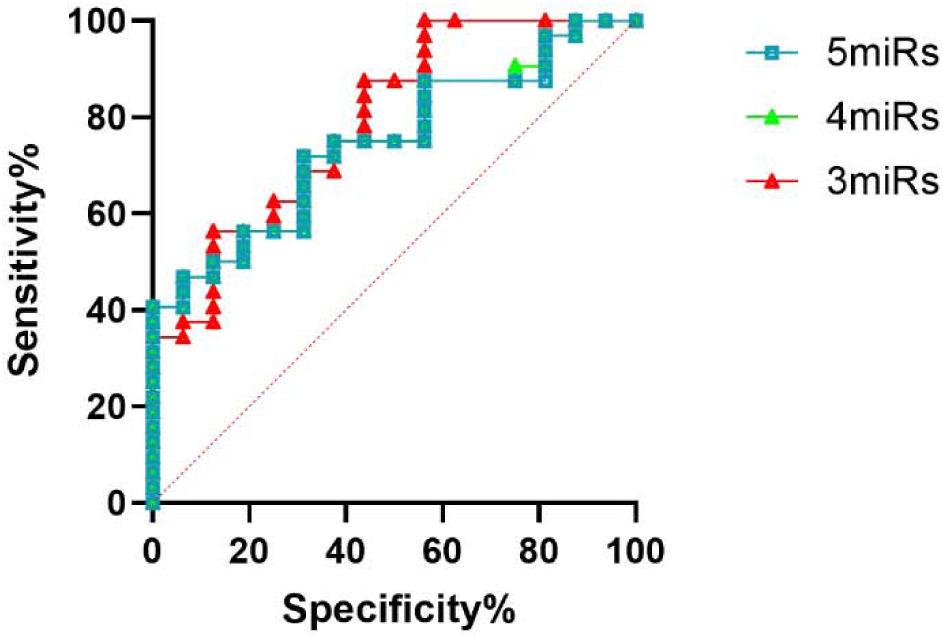
MicroRNAs-6724/7704/10400 and the diagnosis of Urinary Bladder Cancer. Receiver Operator Characteristic Curve of different microRNAs in combination of 5miRs (miR-6724-5p/7704/1273h-5p/200-5p/10400; blue), 4miRs (miR-6724-5p/7704/1273h-5p/200-5p; green), and 3miRs (miR-6724/7704/10400; red), for the detection of UBC; x-axis representing the specificity (%) and y-axis representing the sensitivity (%).

miR-548ae exhibited approximately balanced performance (specificity 57.14%, sensitivity 53.13%, AUC 55.13%), and miR-1273h-5p presented moderate sensitivity and specificity (83.87%, 40%) with an intermediate AUC (61.94%). Collectively, these findings indicate miR-6724-5p and miR-10400-5p as standout candidates with optimal combined sensitivity and AUC.

Based on the AUC values and sensitivity metrics, the most effective three-miRNA panel for UBC biomarker development consists of miR-6724-5p, miR-10400-5p, and miR-7704. This combination maximizes average AUC while maintaining high sensitivity for disease detection. These miRNAs collectively offer superior diagnostic accuracy and are suitable for inclusion as principal candidates in a non-invasive biomarker panel for urinary bladder cancer.

## Discussion

This study delineates urinary mature microRNAs (miRNAs) as highly robust and biologically significant biomarkers for urinary bladder cancer (UBC). The exceptional ex vivo stability of urinary miRNAs under physiological conditions, attributed to their small size, lack of polyadenylation, and protective encapsulation within exosomes, supports their reliable quantification and reproducibility across clinical sample processing workflows (Guhaniyogi & Brewer, 2001; Kural et al., 2024; Mitchell et al., 2008; Valadi et al., 2007). This molecular resilience underpins their growing utility as non-invasive indicators of bladder cancer presence and progression.

High-throughput sequencing revealed a notably expanded repertoire of 865 known and 11 novel miRNAs in urinary samples, with UBC patients displaying a markedly more complex miRNA landscape (708 known miRNAs) than controls (540 known). Differential expression analysis identified 10 miRNAs significantly dysregulated in UBC (p < 0.05, |log2Foldchange| ≥ 1.0), confirming profound cancer-associated remodeling of the urinary miRNA milieu (Braicu et al., 2015; Grimaldi et al., 2022). Given that precursor RNAs lack biological activity (Gogakos et al., 2017; Yang et al., 2019), a stringent computational filtering was employed to prioritize mature miRNAs for biomarker discovery and clinical correlation (Figure 1D, Supplementary Table 1). This approach circumvents technical artefacts and ensures focus on sequence-unique, disease-relevant signatures, in line with current recommendations for miRNA biomarker assay development (Condrat et al., 2020). Among mature miRNAs, four miRNAs; miR-200a-5p, miR-10400-5p, miR-1273h-5p, and miR-6724-5p, were uniquely identified in UBC patients, whereas miR-548ae-5p and miR-7704 were confined to controls. MiR-934, while present in both cohorts, displayed significant upregulation in UBC cases (Supplementary Table 1).

Validation in an age-matched cohort (n=45) demonstrated heterogeneity yet clear disease-linked miRNA expression patterns (Supplementary Table 2 and 3) revealing substantial overexpression of miR-6724-5p (fold change ∼1,494; p=0.0008), miR-1273h-5p (∼1,086; p=0.0069), miR-200-5p (∼2,777; p=0.0011), and miR-10400-5p (∼46; p=0.0011) in cancer samples. The discordance observed in miR-7704 expression across sequencing and qPCR platforms emphasizes the necessity for cross-validation and highlights potential biological complexity or technical variability intrinsic to miRNA detection methodologies.

Stage-specific analyses revealed nuanced expression dynamics: miR-6724-5p peaked in metastatic disease, implicating it as a biomarker of tumor aggressiveness; miR-1273h-5p and miR-7704 were elevated in high-risk and muscle-invasive stages, indicating roles in progression; miR-200-5p, a known oncogenic miRNA (Jo et al., 2022; Klicka et al., 2022), sustained elevated expression across early and intermediate risk cancer but decreased in metastasis, reflecting shifting molecular regulation during disease evolution. These findings advocate for the deployment of multiplexed, disease stage-informed miRNA panels for precision diagnostics.

Receiver operating characteristic (ROC) analyses highlighted miR-6724-5p and miR-10400-5p as top-performing diagnostic markers with AUC values exceeding 70% and sensitivities above 90%. MiR-7704 and miR-200-5p added value through robust sensitivity despite moderate specificity. The combined three-miRNA panel (miR-6724-5p, miR-10400-5p, miR-7704) yielded superior diagnostic accuracy compared to individual markers, in accordance with recent meta-analyses endorsing multiplex urinary miRNA panels as hallmark tools for UBC detection (Li et al., 2022).

In conclusion, this study corroborates urinary mature miRNAs, particularly miR-6724-5p, miR-10400-5p, and miR-7704, as effective non-invasive biomarkers for UBC. Their remarkable stability, reproducibility, and stage-specific expression patterns underscore their translational potential for incorporation into clinical diagnostic and monitoring platforms. However, this study was limited by its small sample size, a shorter follow-up period and its cross-sectional design which precluded the correlation between the identified biomarker miRNAs with cancer recurrence, and patient survival. Future work should aim to validate these findings prospectively across multicenter, larger cohorts and refine assay standardization for clinical deployment.

## Declarations

### Ethical Approval and Consent for participation and publication

The study was approved by the Ethical Committee, IMS-BHU (Ref. No. 2024/EC/7064). Written informed consent was obtained from all participants (in both Hindi/ English) after a detailed explanation of the study protocol.

### Data Availability

The datasets presented and analysed was submitted to the NCBI database. With Sequence Read Archive ID: SUB15663449 .and can be found at (…………………). All the data generated and analysed in this study are available from corresponding author on request.

### Authors’ Contribution

Garima Singh and Anil Kumar was involved in investigation, methodology, data analysis, writing original draft, review & editing of the manuscript. Neelu Mishra, Shristy Bhattacharjee and Kunal Yadav helped in the execution of experiments, data analysis and was involved in critical reviewing and editing of the manuscript. Yashasvi Singh, Ujwal Kumar, and Sameer Trivedi, have been involved in critical review and editing of manuscript. Lalit Kumar was involved in the project administration, supervision, critical review & editing of the manuscript. Samarendra K Singh was involved in the project administration, conceptualization and designing, supervision, critical review & editing of the manuscript

## Acknowledgement

We wish to thank the School of Biotechnology, Institute of Science, BHU and Department of Urology, IMS-BHU for supporting the research. We thank the staff and patients of the Department of Urology, IMS-BHU. We are also thankful to ISLS for facilitating the qRT-PCR facility. We would also like to acknowledge MedGenome Labs Ltd, Bangalore, Karnataka, India for providing the facility for miRNA sequencing. We would like to extend our gratitude towards Department of Biotechnology (DBT), Govt. of India for funding Shristy Bhattacharjee and Kunal Yadav, Council of Scientific and Industrial Research (CSIR), Govt. of India for funding Neelu Mishra. We also thank Department of Science & Technology-Fund for Improvement of S&T Infrastructure (DST-FIST), Govt. of India for providing the infrastructure fund to the School of Biotechnology, BHU.

## Supplementary Materials

**Supplementary Figure 1.**
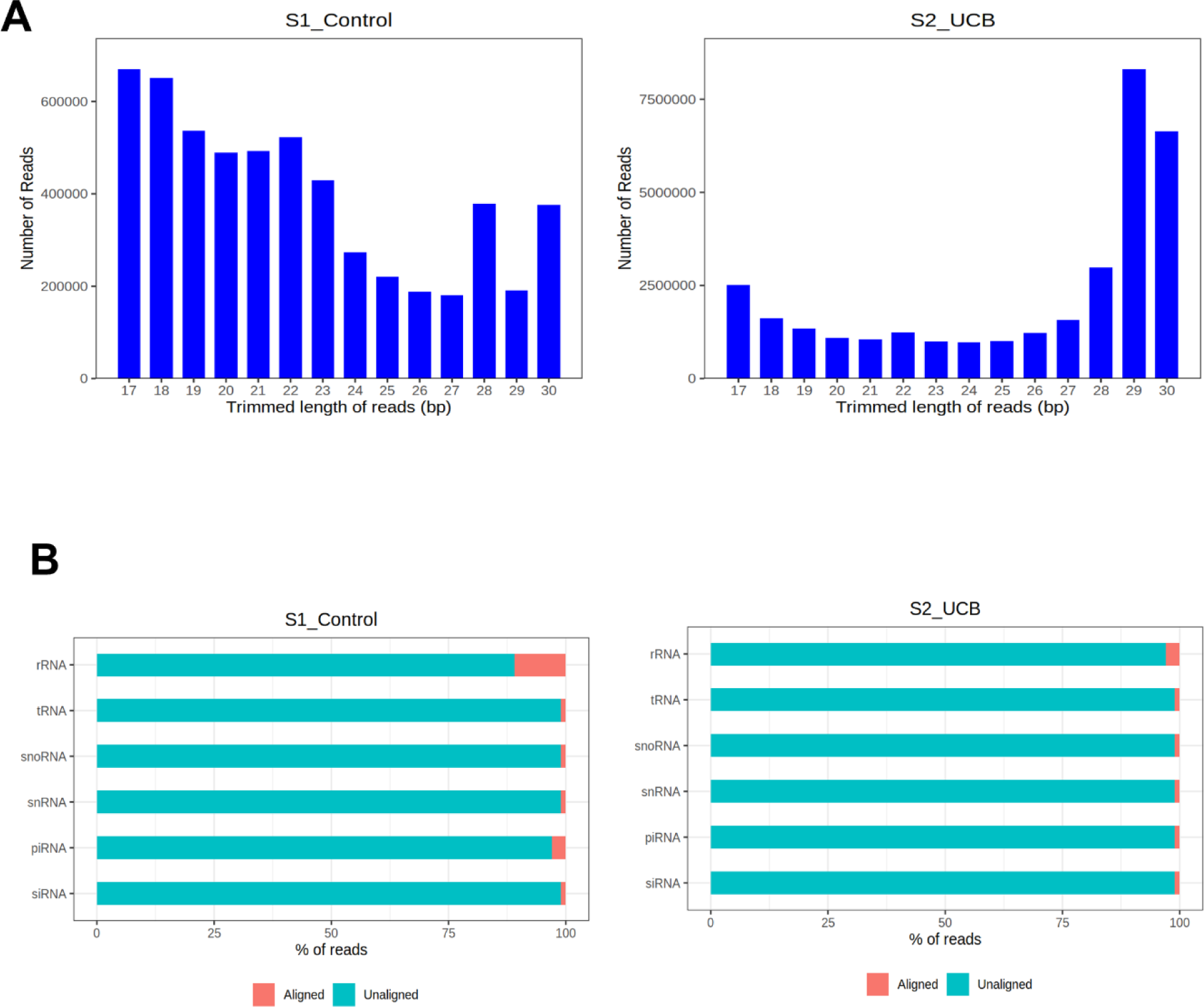
**A.** Number of Reads length distribution for sample S1, Control and S2, Urinary Bladder Cancer. **B.** Small RNA distribution for sample S1, Control and S2, Urinary Bladder Cancer

**Supplementary Table 1:**
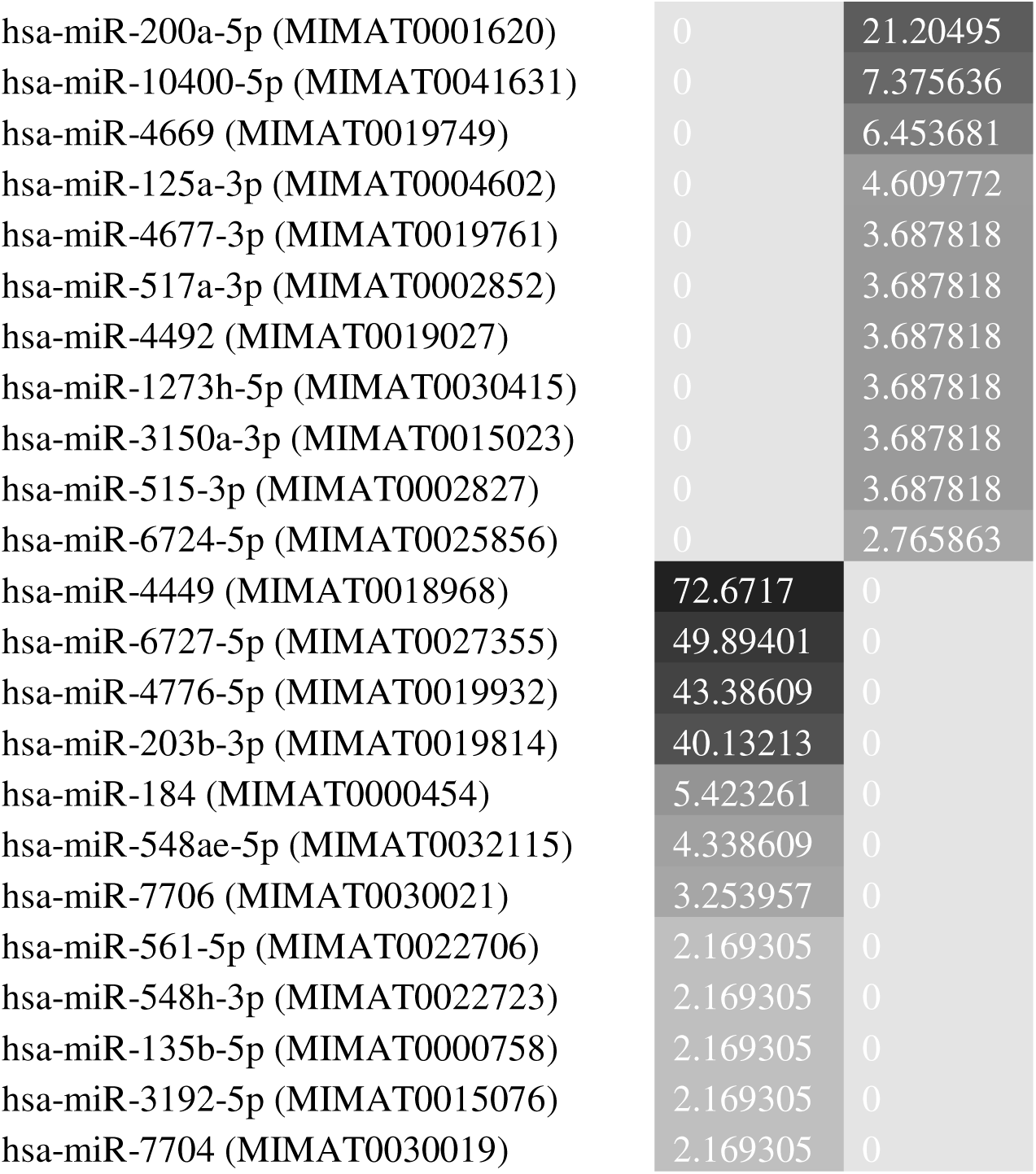
Differential Expression of mature miRNAs in Urine of UBC Patient Sample.

**Supplementary Table 2:**
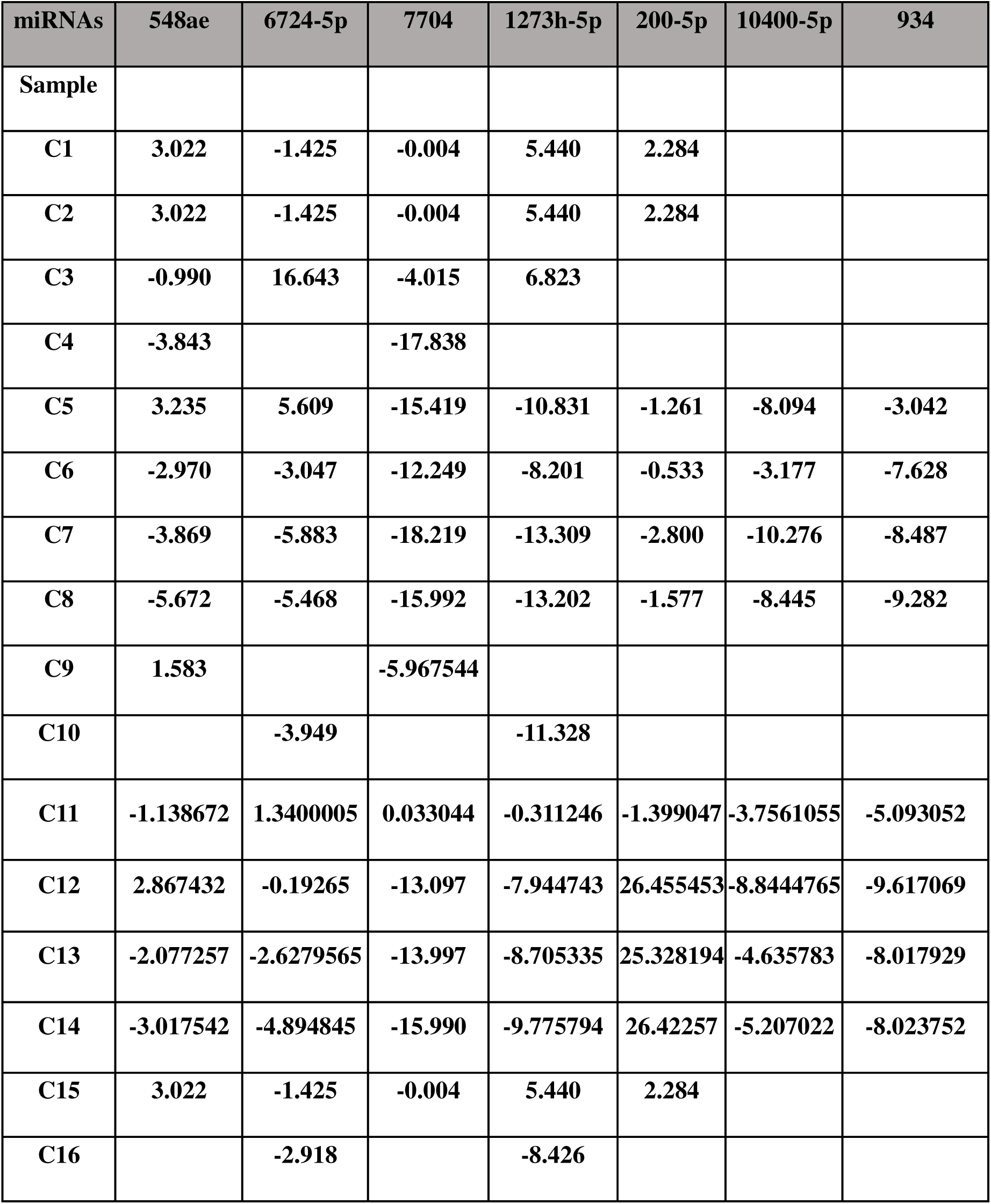
Delta CT for miRNAs in Urine of Control Samples.

**Supplementary Table 3:**
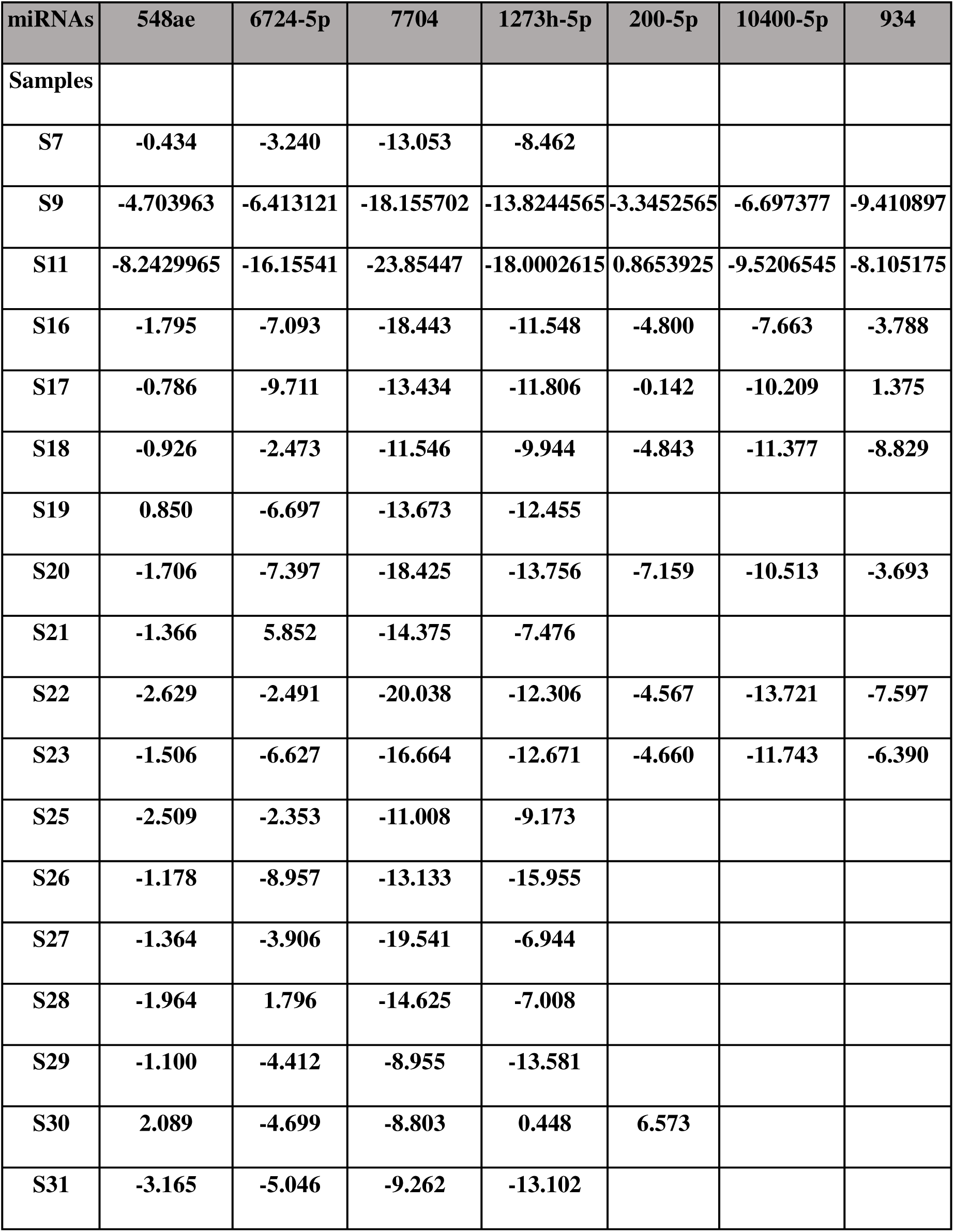
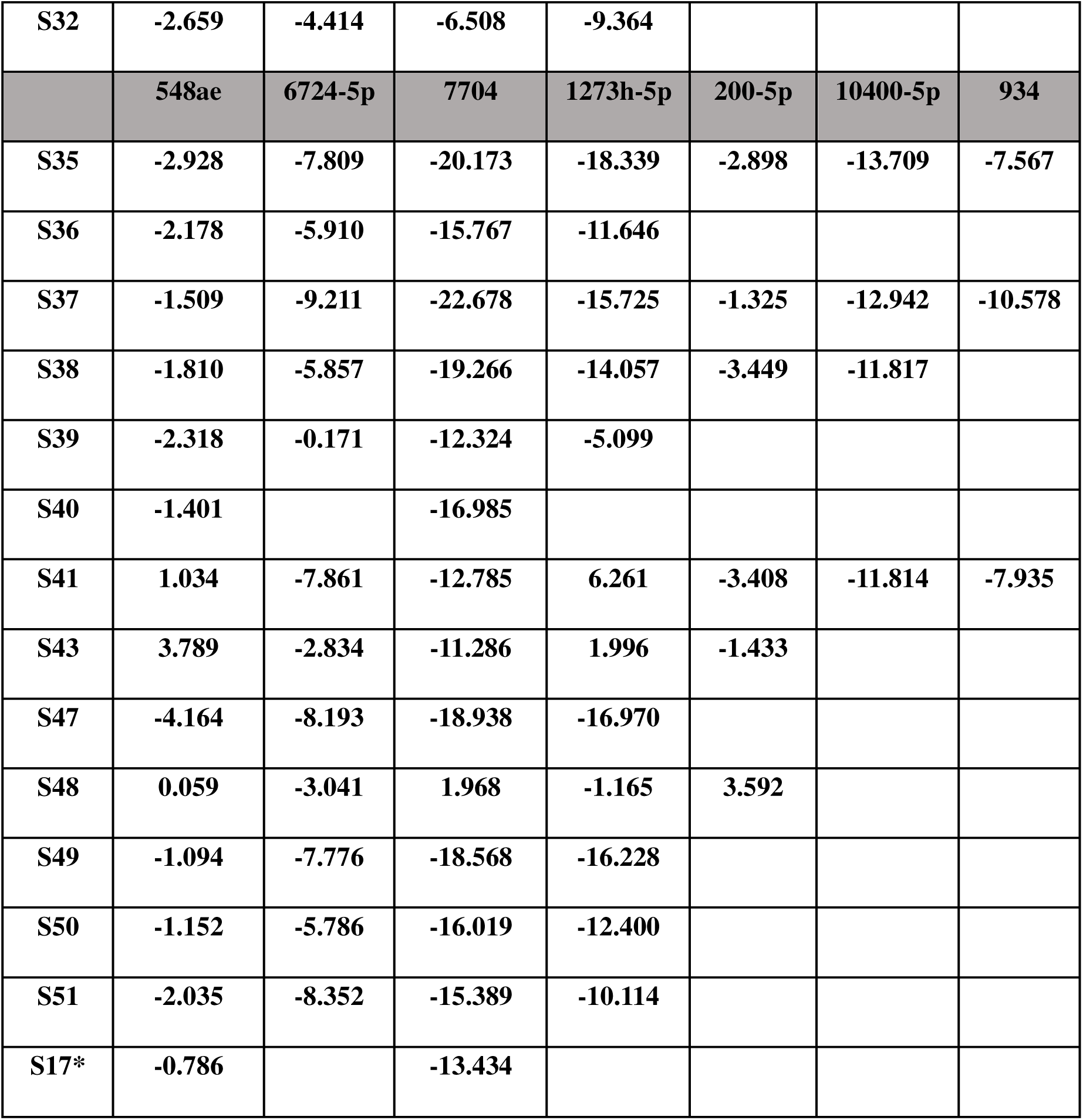
Delta CT for miRNAs in Urine of Urinary Bladder Cancer Patient Samples.

