## Supplementary Figures and Tables for "Urinary Exosomal miRNA Profiling Reveals Sensitive Non-Invasive Diagnosis of Bladder Cancer"

**Supplementary Materials**

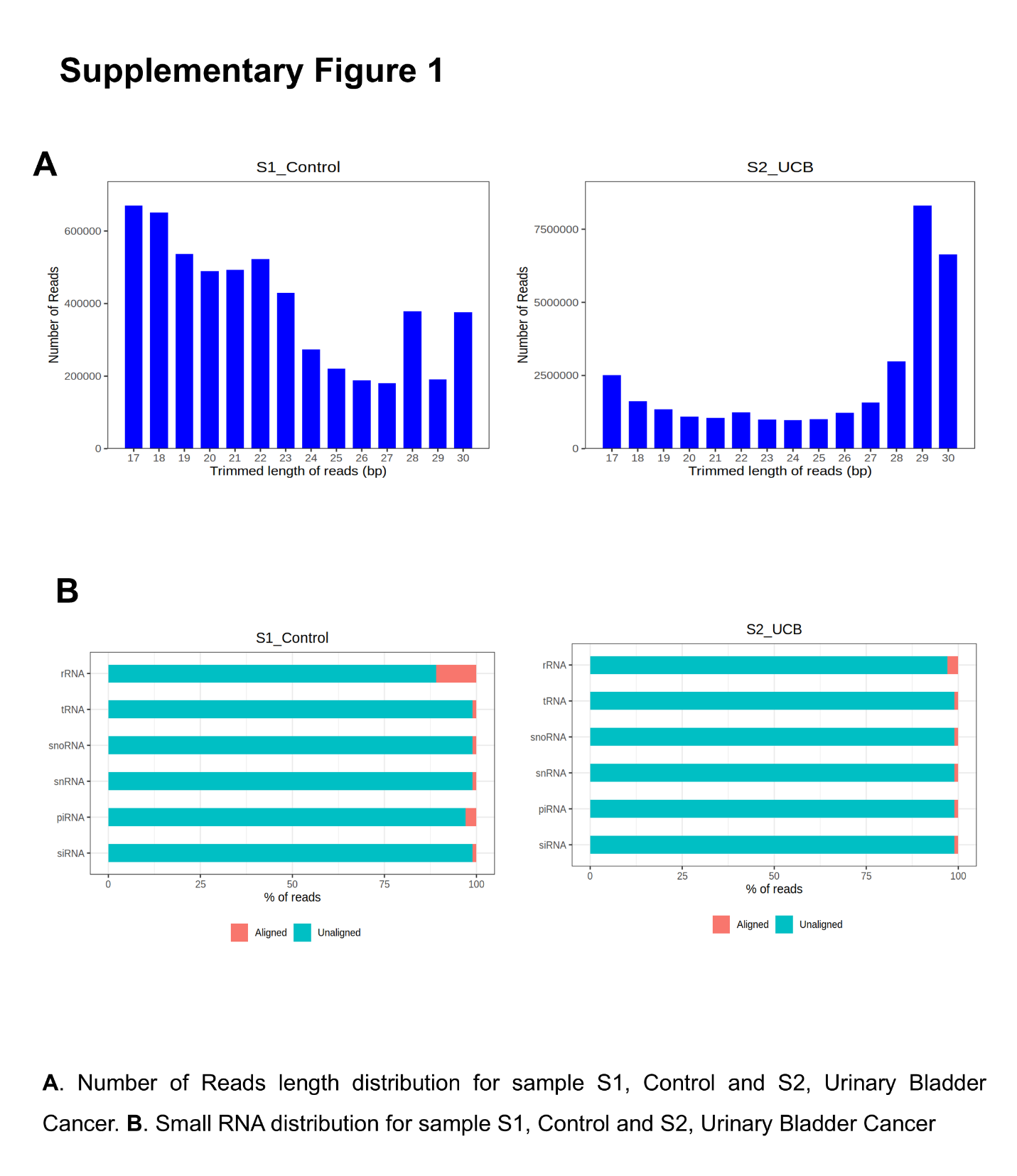

| **Mature miRNA** | **Control** | **UBC** |
| --- | --- | --- |
| hsa-miR-934 (MIMAT0004977) | 1.084652 | 76.52222 |
| hsa-miR-200a-5p (MIMAT0001620) | 0 | 21.20495 |
| hsa-miR-10400-5p (MIMAT0041631) | 0 | 7.375636 |
| hsa-miR-4669 (MIMAT0019749) | 0 | 6.453681 |
| hsa-miR-125a-3p (MIMAT0004602) | 0 | 4.609772 |
| hsa-miR-4677-3p (MIMAT0019761) | 0 | 3.687818 |
| hsa-miR-517a-3p (MIMAT0002852) | 0 | 3.687818 |
| hsa-miR-4492 (MIMAT0019027) | 0 | 3.687818 |
| hsa-miR-1273h-5p (MIMAT0030415) | 0 | 3.687818 |
| hsa-miR-3150a-3p (MIMAT0015023) | 0 | 3.687818 |
| hsa-miR-515-3p (MIMAT0002827) | 0 | 3.687818 |
| hsa-miR-6724-5p (MIMAT0025856) | 0 | 2.765863 |
| hsa-miR-4449 (MIMAT0018968) | 72.6717 | 0 |
| hsa-miR-6727-5p (MIMAT0027355) | 49.89401 | 0 |
| hsa-miR-4776-5p (MIMAT0019932) | 43.38609 | 0 |
| hsa-miR-203b-3p (MIMAT0019814) | 40.13213 | 0 |
| hsa-miR-184 (MIMAT0000454) | 5.423261 | 0 |
| hsa-miR-548ae-5p (MIMAT0032115) | 4.338609 | 0 |
| hsa-miR-7706 (MIMAT0030021) | 3.253957 | 0 |
| hsa-miR-561-5p (MIMAT0022706) | 2.169305 | 0 |
| hsa-miR-548h-3p (MIMAT0022723) | 2.169305 | 0 |
| hsa-miR-135b-5p (MIMAT0000758) | 2.169305 | 0 |
| hsa-miR-3192-5p (MIMAT0015076) | 2.169305 | 0 |
| hsa-miR-7704 (MIMAT0030019) | 2.169305 | 0 |

**Supplementary Table 1: Differential Expression of mature miRNAs in Urine of UBC Patient Sample**

**Supplementary Table 2: Delta CT for miRNAs in Urine of Control Samples**

| **miRNAs** | **548ae** | **6724-5p** | **7704** | **1273h-5p** | **200-5p** | **10400-5p** | **934** |
| --- | --- | --- | --- | --- | --- | --- | --- |
| **Sample** |  |  |  |  |  |  |  |
| **C1** | **3.022** | **-1.425** | **-0.004** | **5.440** | **2.284** |  |  |
| **C2** | **3.022** | **-1.425** | **-0.004** | **5.440** | **2.284** |  |  |
| **C3** | **-0.990** | **16.643** | **-4.015** | **6.823** |  |  |  |
| **C4** | **-3.843** |  | **-17.838** |  |  |  |  |
| **C5** | **3.235** | **5.609** | **-15.419** | **-10.831** | **-1.261** | **-8.094** | **-3.042** |
| **C6** | **-2.970** | **-3.047** | **-12.249** | **-8.201** | **-0.533** | **-3.177** | **-7.628** |
| **C7** | **-3.869** | **-5.883** | **-18.219** | **-13.309** | **-2.800** | **-10.276** | **-8.487** |
| **C8** | **-5.672** | **-5.468** | **-15.992** | **-13.202** | **-1.577** | **-8.445** | **-9.282** |
| **C9** | **1.583** |  | **-5.967544** |  |  |  |  |
| **C10** |  | **-3.949** |  | **-11.328** |  |  |  |
| **C11** | **-1.138672** | **1.3400005** | **0.033044** | **-0.311246** | **-1.399047** | **-3.7561055** | **-5.093052** |
| **C12** | **2.867432** | **-0.19265** | **-13.097** | **-7.944743** | **26.455453** | **-8.8444765** | **-9.617069** |
| **C13** | **-2.077257** | **-2.6279565** | **-13.997** | **-8.705335** | **25.328194** | **-4.635783** | **-8.017929** |
| **C14** | **-3.017542** | **-4.894845** | **-15.990** | **-9.775794** | **26.42257** | **-5.207022** | **-8.023752** |
| **C15** | **3.022** | **-1.425** | **-0.004** | **5.440** | **2.284** |  |  |
| **C16** |  | **-2.918** |  | **-8.426** |  |  |  |

**Supplementary Table 3: Delta CT for miRNAs in Urine of Urinary Bladder Cancer Patient Samples**

| **miRNAs** | **548ae** | **6724-5p** | **7704** | **1273h-5p** | **200-5p** | **10400-5p** | **934** |
| --- | --- | --- | --- | --- | --- | --- | --- |
| **Samples** |  |  |  |  |  |  |  |
| **S7** | **-0.434** | **-3.240** | **-13.053** | **-8.462** |  |  |  |
| **S9** | **-4.703963** | **-6.413121** | **-18.155702** | **-13.8244565** | **-3.3452565** | **-6.697377** | **-9.410897** |
| **S11** | **-8.2429965** | **-16.15541** | **-23.85447** | **-18.0002615** | **0.8653925** | **-9.5206545** | **-8.105175** |
| **S16** | **-1.795** | **-7.093** | **-18.443** | **-11.548** | **-4.800** | **-7.663** | **-3.788** |
| **S17** | **-0.786** | **-9.711** | **-13.434** | **-11.806** | **-0.142** | **-10.209** | **1.375** |
| **S18** | **-0.926** | **-2.473** | **-11.546** | **-9.944** | **-4.843** | **-11.377** | **-8.829** |
| **S19** | **0.850** | **-6.697** | **-13.673** | **-12.455** |  |  |  |
| **S20** | **-1.706** | **-7.397** | **-18.425** | **-13.756** | **-7.159** | **-10.513** | **-3.693** |
| **S21** | **-1.366** | **5.852** | **-14.375** | **-7.476** |  |  |  |
| **S22** | **-2.629** | **-2.491** | **-20.038** | **-12.306** | **-4.567** | **-13.721** | **-7.597** |
| **S23** | **-1.506** | **-6.627** | **-16.664** | **-12.671** | **-4.660** | **-11.743** | **-6.390** |
| **S25** | **-2.509** | **-2.353** | **-11.008** | **-9.173** |  |  |  |
| **S26** | **-1.178** | **-8.957** | **-13.133** | **-15.955** |  |  |  |
| **S27** | **-1.364** | **-3.906** | **-19.541** | **-6.944** |  |  |  |
| **S28** | **-1.964** | **1.796** | **-14.625** | **-7.008** |  |  |  |
| **S29** | **-1.100** | **-4.412** | **-8.955** | **-13.581** |  |  |  |
| **S30** | **2.089** | **-4.699** | **-8.803** | **0.448** | **6.573** |  |  |
| **S31** | **-3.165** | **-5.046** | **-9.262** | **-13.102** |  |  |  |
| **S32** | **-2.659** | **-4.414** | **-6.508** | **-9.364** |  |  |  |
|  | **548ae** | **6724-5p** | **7704** | **1273h-5p** | **200-5p** | **10400-5p** | **934** |
| **S35** | **-2.928** | **-7.809** | **-20.173** | **-18.339** | **-2.898** | **-13.709** | **-7.567** |
| **S36** | **-2.178** | **-5.910** | **-15.767** | **-11.646** |  |  |  |
| **S37** | **-1.509** | **-9.211** | **-22.678** | **-15.725** | **-1.325** | **-12.942** | **-10.578** |
| **S38** | **-1.810** | **-5.857** | **-19.266** | **-14.057** | **-3.449** | **-11.817** |  |
| **S39** | **-2.318** | **-0.171** | **-12.324** | **-5.099** |  |  |  |
| **S40** | **-1.401** |  | **-16.985** |  |  |  |  |
| **S41** | **1.034** | **-7.861** | **-12.785** | **6.261** | **-3.408** | **-11.814** | **-7.935** |
| **S43** | **3.789** | **-2.834** | **-11.286** | **1.996** | **-1.433** |  |  |
| **S47** | **-4.164** | **-8.193** | **-18.938** | **-16.970** |  |  |  |
| **S48** | **0.059** | **-3.041** | **1.968** | **-1.165** | **3.592** |  |  |
| **S49** | **-1.094** | **-7.776** | **-18.568** | **-16.228** |  |  |  |
| **S50** | **-1.152** | **-5.786** | **-16.019** | **-12.400** |  |  |  |
| **S51** | **-2.035** | **-8.352** | **-15.389** | **-10.114** |  |  |  |
| **S17*** | **-0.786** |  | **-13.434** |  |  |  |  |
